# Non-linear stress-softening of the bacterial cell wall confers cell shape homeostasis

**DOI:** 10.1101/2024.09.03.611099

**Authors:** Paola Bardetti, Felix Barber, Enrique R. Rojas

## Abstract

The bacillus - or rod - is a pervasive cellular morphology among bacteria. Rod-shaped bacteria elongate without widening by reinforcing their cell wall anisotropically, along the cell’s circumference, but it is unknown how cells adaptively tune anisotropy to homeostatically control cell width. Through super-resolution measurements of cell wall mechanical properties, we discovered that the *Bacillus subtilis* cell wall exhibits non-linear stress-softening exclusively in the circumferential direction. Furthermore, during steady-state growth the cell wall is inflated precisely to the acute non-linear transition. Physics-based theory correctly predicted that this transition underlies the negative feedback that governs cell width homeostasis. In other words, the cell wall is a “smart material” whose exotic mechanical properties are exquisitely adapted to execute cellular morphogenesis.

## Introduction

The peptidoglycan cell wall is a covalently cross-linked polymer network that defines the size and shape of bacterial cells (Fig. 1A). Three processes mediate expansion of the wall during cell growth: peptidoglycan synthesis^1^, enzymatic hydrolysis of the peptide moieties^2^, and mechanical deformation of the wall by the large intracellular turgor pressure^3^ (≈1-30 atm, depending on species^4, 5^). The primary protein machinery that executes peptidoglycan synthesis during the growth of rod-shaped bacteria is the Rod complex, a heterocomplex of 5 proteins that is scaffolded by oligomers of bacterial actin-homologues^6-8^. These transmembrane complexes synthesize peptidoglycan processively, resulting in anisotropic cell wall microstructure in which the glycan moieties are oriented approximately circumferentially^9^ (*θ*, Fig. 1A) and are connected to one another via the peptide moieties.

**Figure 1.**
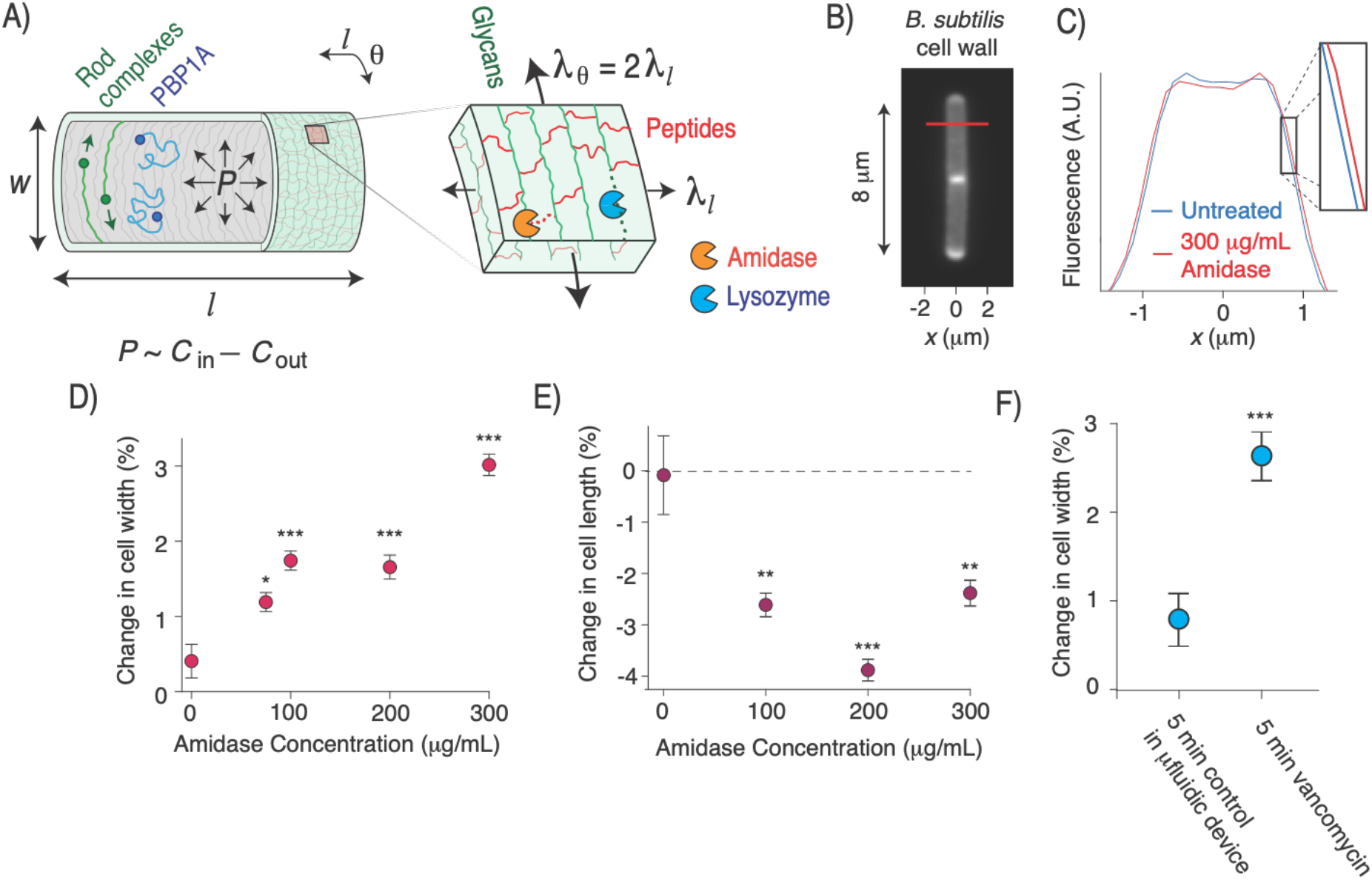
Exogenous peptide digestion causes cell widening. A) Diagram of the Gram-positive cell wall including its modes of synthesis and cell surface co-ordinate system (*l,θ*). *P*: turgor pressure. A_1,θ_ are the longitudinal and circumferential surface tensions. B) *B. subtilis* cell wall labeled with the fluorescent D-amino acid HADA. C) Mean fluorescent profile across the width of the labeled cell wall (red line in B) before and after 30 s amidase treatment. D) Mean change in cell width upon amidase treatment. Error bars indicate +/-1 s.e.m. *n*=15-50 cells across two technical replicates per data point. E) Mean change in cell length upon amidase treatment, controlling for cell growth. Error bars indicate +/-1 s.e.m. *n*=20-50 cells across two technical replicates per data point. F) Change in cell width during 5 minute of growth in the microfluidic chip (control) and during 5 minutes of treatment with 10 mg/mL vancomycin. ^*^: *p*=10^−2^,^**^: 10^−3^,^***^: *p*<10^−4^, compared to untreated control.

It is widely believed that the oriented glycans polymers promote rod-shaped growth by mechanically reinforcing the cell wall in the circumferential direction, effectively “girdling” the cell to prevent turgor pressure from driving cell widening^10^. It is also commonly assumed that peptides are oriented longitudinally^11^ (*l*, Fig. 1A). If so, hydrolysis of load-bearing peptides would tend to cause cellular elongation rather than widening.

However, this model is not sufficient to explain rod-shaped morphogenesis. Glycans are not oriented strictly circumferentially^9^ nor are they infinitely stiff^4^. Similarly, peptides are not oriented strictly longitudinally, and in Gram-positive bacteria the requirement for peptides to connect glycans across the thickness of the cell wall make this impossible. Finally, inflation of the cell by turgor pressure causes anisotropic surface tension in the cell wall whereby the circumferential tension is twice the longitudinal tension^12^ (*λ*_*θ*_ = 2*λ*_*l*_; Fig. 1A). Therefore, based on first principles, hydrolysis of peptide moieties during cell growth is expected to lead to both elongation and widening.

### Peptide hydrolysis causes cell widening

To explicitly demonstrate this principle, we measured the change in cell width in response to exogenous digestion of cell-wall peptides in the Gram-positive bacterium *Bacillus subtilis*. To do so, we used microfluidics to acutely perfuse single bacterial cells with a purified recombinant amidase (Fig. 1A) derived from the lytic bacteriophage SPP1^13^. We labeled the cell wall with a fluorescent D-amino acid^14^ and developed a super-resolution method to quantify nanometer-scale changes in cell width (Fig. 1B,C). As hypothesized, amidase perfusion led to a dose-dependent increase in width (Fig. 1D). Surprisingly, when controlling for cell growth, amidase also caused a decrease in cell length (Fig. 1E, *Methods*). Perfusion of lysozyme, which digests the glycan moieties led to the same qualitative effects (Fig. S1A,B).

As an alternative method to measure the effect of hydrolysis on cell width, we perfused cells with an inhibitory concentration of vancomycin, which prevents cell wall synthesis without completely inhibiting hydrolysis^15^, leading eventually to lysis (Fig. S1C). This treatment also caused cell widening even before it had an effect on elongation rate (Fig. 1F,S1C). Together, these data indicate that peptidoglycan hydrolysis, when unbalanced by synthesis, leads to cell widening.

In light of these data, we hypothesized that to avoid slow increases in cell width during the peptidoglycan hydrolysis required for cell growth, cells exert a radially constrictive force on the cell wall that opposes turgor pressure. It was previously shown computationally that applying pre-tension to nascent glycan polymers could prevent pressure-driven widening^16^, but this mechanism was not tested, nor is there an obvious molecular motor within the Rod complex that could exert tension on the cell wall.

### Bacteria exhibit finger-trap mechanics

To probe the magnitude of the constrictive force that would be required to counteract inflation, we first measured the mechanical properties of the cell wall. Atomic force microscopy is commonly used to measure the deformation of the cell wall in response to indentation forces^4^, but the wall’s deformation with respect to changes in surface tension is more relevant to cellular morphogenesis. We therefore subjected *B. subtilis* cells to a microfluidics-based “osmotic force-extension” assay that we previously developed for Gram-negative bacteria^17^. In this assay, single cells are subjected to a series of osmotic shocks (Fig. 2A), which cause acute changes in pressure. These pressure variations deform the cell wall (Fig. 2B), and the dependence of this deformation (the longitudinal and circumferential strains, *ε*_*l*_ and *ε*_*θ*_) on shock magnitude yields an empirical measurement of the anisotropic mechanical properties of the cell wall (Fig. 2C,D,E).

**Figure 2.**
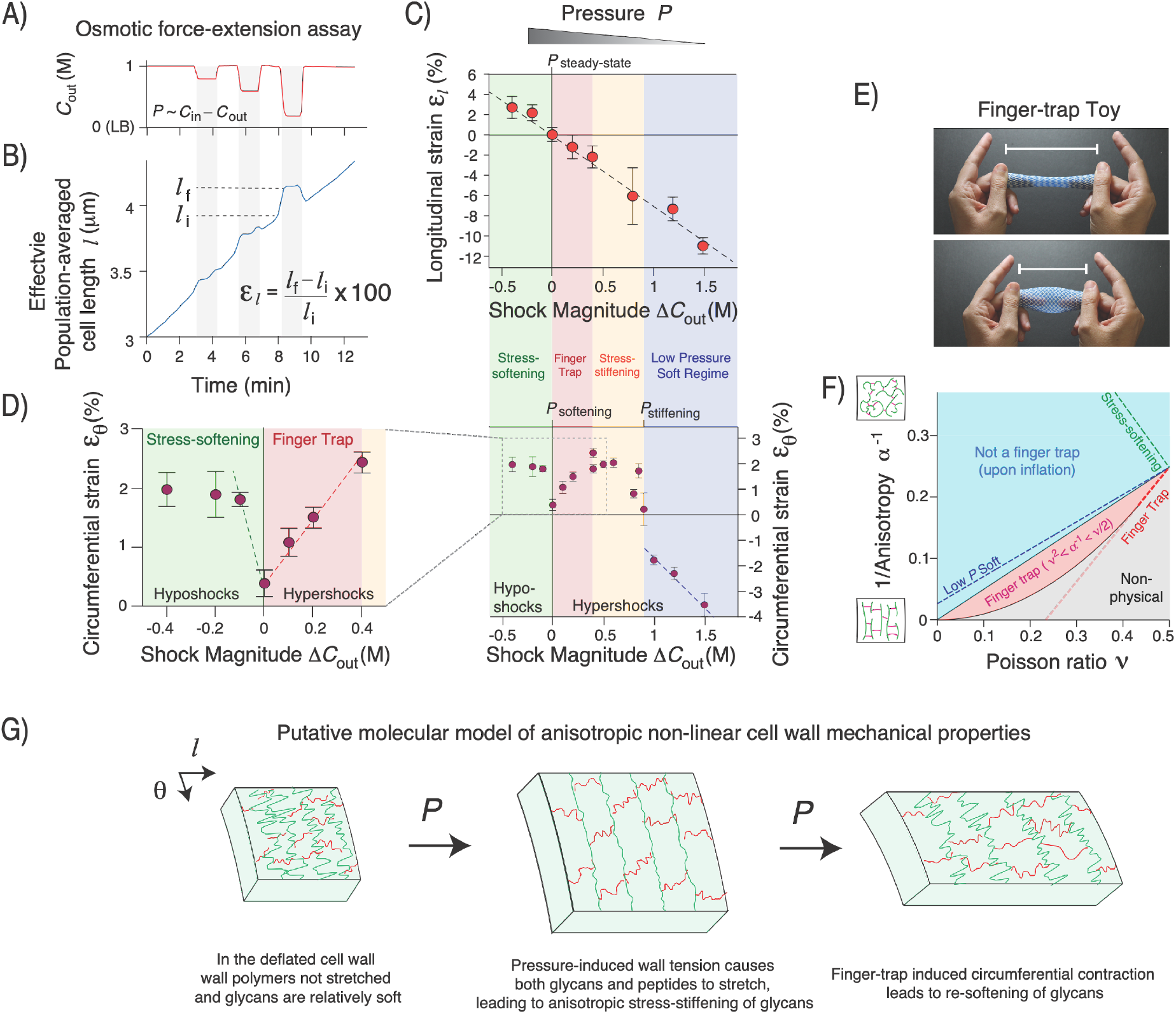
The cell wall exhibits anisotropic stress-stiffening and stress-softening. A) Extracellular concentration of sorbitol, *C*_out_, versus time during an osmotic force-extension experiment. B) Effective population-averaged cell length versus time during the osmotic force-extension experiment. C) Longitudinal strain versus shock magnitude across several osmotic force extension experiments. n=20-60 cells across 1-3 replicate experiments per shock magnitude. Error bars: +/-1 s.e.m. The dotted line is a linear regression. D) Circumferential strain versus shock magnitude across several osmotic force extension experiments. n=74-174 across 1-2 replicate experiments per shock magnitude. Error bars: +/-1 s.e.m. The dotted lines are linear regressions to the three linear regimes of mechanical behavior. E) A finger trap toy. F) The mechanical behavior of an inflated linear elastic cylindrical cell wall versus its anisotropy and Poisson ratio. The dotted lines are the behavior constrained by slopes of the regressions in C) and D). G) Putative qualitative molecular model of the anisotropic non-linear cell wall mechanical properties.

As expected, hyperosmotic shocks, which decrease pressure, caused longitudinal contraction of the cell wall (negative longitudinal strain, *ε*_*l*_ < 0), whereas hypoosmotic shocks caused longitudinal stretching (*ε*_*l*_ > 0). Moreover, longitudinal strain was inversely proportional to shock magnitude and reversible (Fig. 2B,C), demonstrating that the cell wall is linearly elastic in this dimension.

As expected, small hypoosmotic shocks, which increase turgor pressure, caused circumferential stretching of the cell wall (Fig. 2C). Counterintuitively, small hyperosmotic shocks also caused circumferential stretching. As a result, in this regime variation in pressure had the opposite effect on cell length and width, similar to the effect of peptidoglycan hydrolysis (Fig. 1D,E,S1A,B). This behavior is reminiscent of a finger-trap toy in which longitudinal tension causes circumferential constriction due to mechanical coupling between the two directions (Fig. 2D). Finger-trap mechanics are more surprising in the case of a pressurized cell where the circumferential surface tension is twice that of the longitudinal tension (Fig. 1A).

Bacterial cells exhibited linear finger-trap behavior for hyperosmotic shocks up to 400 mM (Fig 2C), however, we observed dramatic non-linear behavior outside this range. First, for hyperosmotic shocks between 400 − 800 mM, the circumferential strain plateaued. At ≈800 mM, we observed a sharp decrease in strain, whereas for larger shocks circumferential strain decreased linearly. Second, since small hypoosmotic shocks caused swelling of cell width, steady-state growth occurs precisely at a sharp non-linear transition.

### The cell wall exhibits anisotropic stress-stiffening and -softening

Because the dependence of mechanical strains were approximately linear within the low-pressure and finger-trap mechanical regimes, it was instructive to use linear elasticity theory to interpret these data. For an anisotropic two-dimensional material, linear elasticity can be expressed:

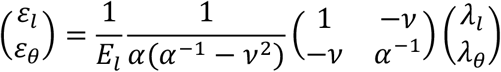

where *E*_*l*_ is the longitudinal elastic modulus (in 2D), *α* = *E*_*θ*_/*E*_*l*_ is the mechanical anisotropy (ratio of the principal elastic moduli), and *ν* is similar to the Poisson ratio (*Methods*). The surface tensions, *λ*_*l*_ = *PR*/2 and *λ*_*θ*_ = *PR*, balance pressure^12^. Imposing the condition that *ε*_*l*_ and *ε*_*θ*_ have opposite signs when pressure is altered reveals that the cell behaves as a finger trap if *α*^−1^ < *ν*/2, that is, if the cell wall is sufficiently anisotropic (Fig. 2F).

Although the empirical dependence of the strains on osmotic shock magnitude (Fig. 2C,D) does not uniquely define *α* and *ν*, it constrains them to a slice of parameter space (Fig. 2F). In the finger-trap regime, this analysis demonstrates that the effective Poisson ratio is between 0.45 < *ν* < 0.5 and *α* is as high as is physically possible within that range, assuming there is no other source of tension other than pressure.

A similar analysis for the mechanical behavior in the low-pressure regime demonstrates that the cell wall has a lower value of anisotropy than in the finger trap regime (Fig. 2F). Since the linear dependence of the longitudinal strain is independent of pressure (Fig. 2C), this means that the transition from the low-pressure regime to the finger-trap regime reflects anisotropic stress-stiffening^4^ exclusively in the circumferential direction.

The observation that cells do not behave as a finger trap for hypoosmotic shocks indicate that the wall is softening in the circumferential direction when pressure is increased beyond its steady-state value. We estimated the degree of softening for small increases in pressure, which revealed that the cell wall is quantitatively much softer in this regime than it is in the finger trap regime or the low-pressure regime (Fig. 2F). Therefore, the cell wall undergoes anisotropic (circumferential) stress-stiffening and stress-softening at different critical values of pressure (Fig. 2C).

These data suggest a qualitative model of *B. subtilis* cell wall mechanics (Fig. 2G). First, at low pressures the cell wall is relatively soft in the circumferential direction because the glycan moieties are not fully extended (circumferential strain is low). For example, the glycans may be in folded conformations. In this soft state, the glycans can be thought of as circumferential slack. Therefore, increasing pressure causes the cell wall to stretch circumferentially, and causes the cell to widen. However, eventually the glycans undergo conventional stress-stiffening (they become taut), causing the wall to stiffen in the circumferential direction and leading to finger-trap mechanics. In this state, as pressure increases the cell wall contracts circumferentially, decreasing cell width. Finally, this contraction eventually causes glycans to exit their taut state, leading to softening of the cell wall in the circumferential direction and loss of finger-trap mechanics. Further increase of pressure causes cell widening. In this model, not only does finger-trap behavior inherently depend on mechanical coupling between the longitudinal and circumferential coupling, but the circumferential stiffening and softening must also depend on longitudinal strain since otherwise as soon as the finger trap was engaged it would contract the wall out of the finger-trap regime.

### Finger-trap mechanics are correlated with cell-width control

Because finger-trap mechanics correspond to constriction of cell width upon elongation, our results raised the possibility that this behavior was required to avoid runaway widening during cell growth. To explore this hypothesis theoretically, we first constructed a simple physical model for the irreversible mechanical expansion of a linear elastic layered cell wall that accounts for variation of cell length and width (Fig. 3A). The specific spatial relationship of peptidoglycan synthesis and hydrolysis during cell growth is unknown, but since rod complexes are membrane-bound it is believed that synthesis of non-load-bearing peptidoglycan occurs proximal to the membrane whereas the hydrolysis of load-bearing peptidoglycan required for cell growth occurs distally^18^. We therefore considered a model in which anisotropic peptidoglycan is added to the inner face of the cell wall at a constant rate and removed at the same rate from the outer face via hydrolysis (*Methods*). Since the cell wall is under tension, removal of load-bearing peptidoglycan dissipates energy and causes irreversible deformation, or growth.

**Figure 3.**
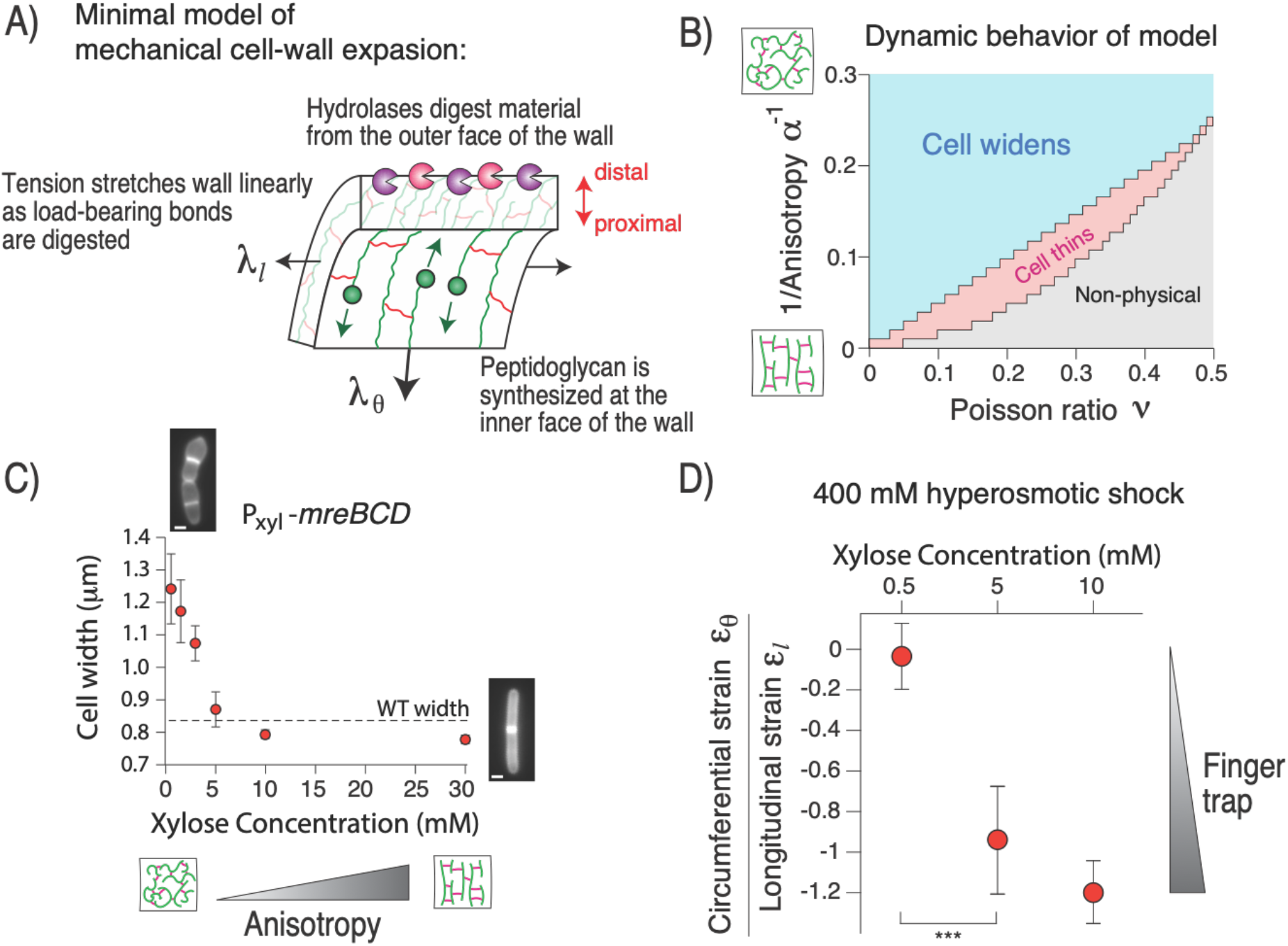
Finger trap mechanics are correlated with cell-width control. A) Diagram of the components of the theoretical linear elastic model for cell growth. B) Dynamic behavior of the model versus the anisotropy and Poisson ratio. C) Cell width versus induction of *mreBCD*. n=22-57 cells across 1 replicate per induction level. Error bars: +/-1 s.d. D) Ratio of circumferential strain and longitudinal strain upon 400 mM hyperosmotic shock for three induction levels of *mreBCD*. Values were calculated from n=26-90 cells across 1-3 replicate experiments (circumferential strain) and n=60-78 cells across 1 replicate experiment (longitudinal strain) per induction level. Error was propagated from the s.e.m. for each respective measurement. ^***^: *p*<10^−4^.

When we solved this model computationally across *α* and *ν*, we found that the region of parameter space in which cell width decreases dynamically was identical to the region where the cell wall behaves as a finger trap mechanically (compare Fig. 3B to 2F). Similarly, in the region of parameter space where cell width increases upon inflation, the cell widens during simulations of cell growth. The reason for the correspondence between the elastic behavior and growth-dynamics is simple: when load-bearing peptidoglycan is removed from the cell wall, the wall must deform in order to balance pressure, and since we analyzed a linear model, the cell wall deforms as a finger trap regardless of the strain distribution along the proximo-distal axis (Fig. 3A).

Therefore, our model made the strong prediction that the cell wall must behave as a finger trap, mechanically, in order to maintain rod shape during growth. We tested this explicitly by genetically manipulating cell-wall mechanical properties and measuring the effect on cell width. It is not possible to tune the Poisson ratio (*ν*) independently, the anisotropy of the cell wall can be systematically altered by titrating components of Rod complex (specifically the *mreBCD* operon) via inducible expression^19^. When biosynthetic flux through the Rod complex is reduced, the balance is accommodated by PBP1, a non-essential enzyme that synthesizes peptidoglycan isotropically (Fig. 1A). Therefore, when the relative expression of the Rod complex is reduced beyond wild-type levels, cell width acutely increases and cells partially lose rod shape (Fig. 3C).

As our model predicted, for Rod complex expression high enough to generate well-shaped rods, modest hyperosmotic shock led to circumferential stretching (finger trap mechanics; Fig. 3D). Conversely, when Rod complex expression was reduced below this threshold cells began to lose rod shape and no longer stretched circumferentially upon hyperosmotic shock. That is, there was a precise correlation between whether the cell maintained a rod shape and whether the cell exhibited finger-trap mechanics. Importantly, for an induction level of 0.5 mM, well below that required for rod-shaped growth, linear elasticity analysis reveals that the cell wall is still highly anisotropic (*α*>4), demonstrating that finger-trap mechanics, rather than anisotropy alone, is the critical factor underlying the mechanics of rod-shaped morphogenesis.

### Pressure-induced wall softening confers width homeostasis

Our analysis explained how cells avoid inflationary widening during cell growth, but the dynamics of this model are inherently unstable: if the cell wall is not a finger trap then the cell widens *ad infinitum* during growth, whereas if the wall is a finger trap then cell width asymptotically approaches zero during growth (Fig. 3B).

However, the linear model ignored the non-linear stress-softening transition (Fig. 2D). Conspicuously, during steady-state growth the cell wall is inflated precisely to this transition. Given this observation, we posited that the transition could confer an intrinsic mechanism for homeostatic control of cell width if there was feedback between cell width, pressure, and/or the mechanical properties of the cell wall (e.g. the critical pressure at which the transition occurs). More specifically, we hypothesized that during steady-state growth cell width is constant because cells are inflated to the non-linear transition, where neither the finger trap nor stress-softening dominate width dynamics. Accordingly, we predicted that for cells that are wider than their steady-steady width but are dynamically thinning (i.e., relaxing to steady-state, Fig. S2), that the cell wall would be in the finger trap regime.

To test this, we cultured cells to steady-state growth at low Rod-complex expression, acutely induced high Rod-complex expression, and then performed circumferential osmotic-force-extension experiments before, during, and after relaxation to steady-state. As for wild-type bacteria, both hypo- and hyperosmotic shocks caused swelling of width for cells grown to steady-state at low, medium, and high Rod complex expression (Fig. 4A-C). However, as we hypothesized, hypoosmotic shocks up to 600 mM caused constriction of cell width for cells that were actively thinning (Fig. 4C). In other words, during cell thinning the cell wall was squarely in the finger trap regime. The validation of this non-intuitive prediction strongly supports our model that the non-linear properties of the cell wall govern width dynamics.

**Figure 4.**
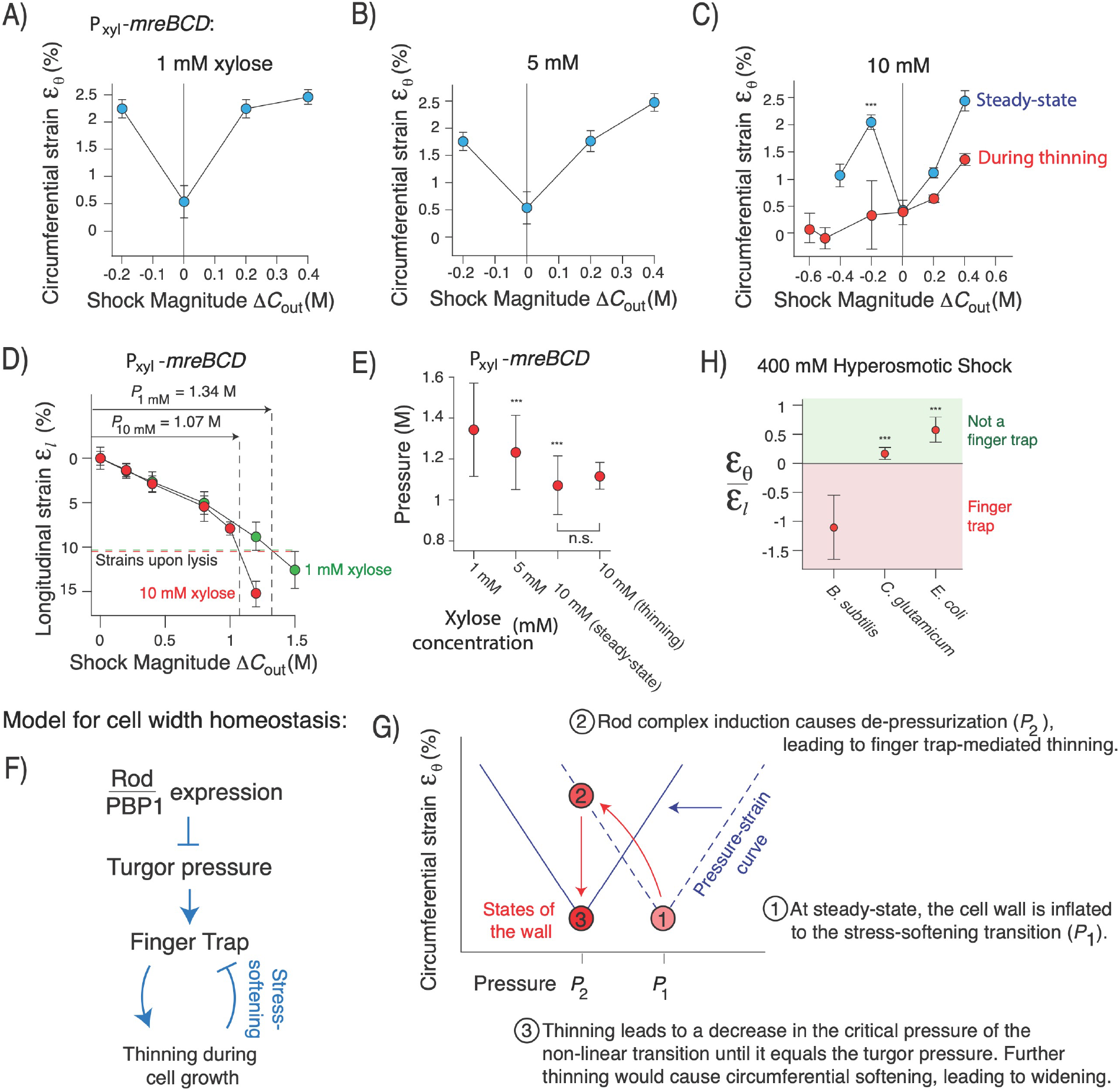
Strain-softening confers cell width homeostasis. Circumferential strain versus shock magnitude at A) 1 mM, B) 5 mM and C) 10 mM xylose induction of P_xyl_-*mreBCB*. n=30-50 cells across 1-3 replicate experiments per shock magnitude. Error bars: +/-1 s.e.m. D) Assay to measure turgor pressure by measuring the magnitude of hyperosmotic shock that causes the cell wall to contract to its rest length. n=60-175 cells across 1-2 replicate experiments per shock magnitude. n=73-90 cells across 2 replicate experiments for each lysis strain measurements. E) Mean turgor pressure versus induction of P_xyl_-*mreBCB* across replicate experiments per induction level. Error was propagated from that of the lysis strain and regression of the osmotic force-extension (*Methods*) F,G) Qualitative negative feedback model of cell width homeostasis based on finger-trap mechanics and stress-softening. H) Ratio of circumferential strain and longitudinal strain upon 400 mM hyperosmotic shock for wild-type *B. subtilis, C. glutamicum*, and *E. coli. B. subtilis* statistics are the same as in Fig. 2C,D. For *C*.*g*.: n=20-30 cells across 1 experiment for circumferential strain and n= 306 across 1 experimental for longitudinal strain. For *E*.*c*.: n=22-37 cells across two experimental replicates for circumferential strain n=238 cells across 3 experimental replicates for longitudinal strain. ^***^: *p*<10^−4^, compared to *B. subtilis*.

Our observation that the cell wall is in the finger trap regime during thinning but not during steady-state growth suggested that turgor pressure is variable during width dynamics. In this light, we next interrogated the nature of the feedback that underlies the adaptation of cell wall mechanical properties during width homeostasis. Since finger trap-mediated cell thinning occurs when turgor pressure is lower than the critical stress-softening pressure, we considered two hypotheses for the mechanism of cell width homeostasis: i) that thinning leads to increased pressure, or ii) thinning leads to a decrease of in the critical stress-softening pressure. Either of these processes would complete a negative feedback loop.

To measure pressure, we performed our longitudinal osmotic-force-extension assay (Fig. 2C), and in a second experiment we lysed cells and measured length contraction upon loss of turgor pressure; we thus quantified the magnitude of hyperosmotic shock that caused contraction of the cell wall to the same length as cell lysis, which gives an empirical measurement of turgor pressure in units of osmolarity (Fig. 4D).

Contrary to our initial expectations, we found that turgor pressure was inversely dependent on Rod complex expression (Fig. 4E). This corresponded to a positive correlation between pressure and cell width during steady-state growth. Since, at steady-state, cells are inflated to the non-linear stress-softening transition regardless of Rod complex expression level, these measurements led us to a surprising conclusion: that the critical stress-softening pressure is dynamically variable.

Despite the correlation between pressure and cell width during steady-state growth, upon acute induction of Rod complex expression pressure equilibrated to its low, steady-state value faster than cell width or the critical pressure of the non-linear transition (Fig. 4C,E), implying that pressure is directly dependent on the specific molecular architecture of the cell wall rather than directly on width. In concurrent work, we identified a molecular mechanism by which *B. subtilis* directly senses cell wall microstructure and relays this information to the osmoregulation systems via cyclic-di-AMP signaling, which could underlie these observations.

Together, our data are consistent with a hybrid feedback system that incorporates molecular and mechanical signals to ensure cell width homeostasis (Fig. 4F,G). According to this model, both turgor pressure and the critical pressure at which the non-linear stress-softening transition are key dynamical variables. If cells are wider than their target steady-state value, then the critical pressure is higher than the actual pressure, the cell wall is in the finger trap regime, and cells thin dynamically. According to our model, thinning causes the critical pressure to decrease until it reaches the actual pressure; further thinning would cause the critical pressure to decrease beyond the actual pressure, positioning the cell wall in the stress-softened regime, which would cause them to widen, completing the negative feedback.

### Finger-trap mechanics are unique to Gram-positive rod-shaped bacteria

We tested the generality of the finger-trap mechanism of cell width control across other bacteria. Interestingly, *Corynebacterium glutamicum*, another rod-shaped Gram-positive bacterium does not exhibit finger trap mechanics for hyperosmotic shocks. This makes sense, however, since *C. glutamicum* elongates via polar growth rather than the diffuse growth used by *B. subtilis*: for polar growth, cell width must be maintained through control of the size of the polar growth zone. Previous measurements of *E. coli* cell geometry in response to osmotic shocks did not indicate finger-trap mechanics^20^, a result we confirmed (Fig. 4H). This suggests that Gram-negative bacteria must also use another mechanism to control width in spite of turgor pressure, which is however much lower than in Gram-positive bacteria^4^.

## Discussion

Transduction of forces within eukaryotic cells is mediated by molecular mechanosensors^21^ that elicit biochemical and genetic responses that can, in turn, manipulate forces within the cell. For example, when force is applied to branched actin networks, the network rapidly increases the amount of force it can sustain through structural alterations to the network, a process that is hard-wired into the biochemistry of actin-binding proteins^22^. Like actin, we found that the Gram-positive bacterial cell wall is a “smart material” that internally senses and responds to forces. However, the cell wall employs a quintessentially prokaryotic paradigm whereby negative-feedback regulation of cell width is embedded the exotic non-linear properties of the wall.

A major open question motivated by our discovery is the specific mechanism by which the critical pressure at which the non-linear transition occurs depends on cell width. We hypothesize that a key variable that is missing from our model is the surface area-to-mass ratio, which determines the metabolic flux through the peptidoglycan biosynthesis pathway^15^. For example, the critical pressure could depend on the quantitative ratio between this flux and that of peptidoglycan hydrolysis, which is less likely to depend on cell width. Building upon our analysis, such a mechanism would provide scaling relationships that predict how cell width quantitatively depends on environmental variables including nutrients and osmotic pressure.

## Acknowledgements

We thank Ethan Garner and Carlos São-José for strains, plasmids, technical assistance, and helpful discussions.

## Supplementary Materials

Materials and Methods

Figs. S1 and S2

Table S1

