## Supplementary Information for "Non-linear stress-softening of the bacterial cell wall confers cell shape homeostasis"

### Materials and Methods

**Bacterial strains and growth condition**. The strains used in this study are listed in Table S1. Cells were grown in Luria-Bertani (LB) medium at 37°C with continuous shaking. For promoter induction of the strain *Pxyl-mreBCD*, cells cultured overnight in LB supplemented with 3mM xylose, were diluted in fresh media with the indicated amount of xylose until exponential phase was reached. Hypoosmotic shocks were done by using media supplemented with the indicated amount of sorbitol; for hypoosmotic shocks, cells from overnight culture were diluted in LB medium supplemented with 1M sorbitol; lower concentration of sorbitol were used to shock the cells. *Corynaebacterium glutamicum* cells were grown at 30°C in Brain Heart Infusion medium (BHI); to perform osmotic shocks BHI media was supplemented with the indicated amount of sorbitol.

**Microscopy in microfluidic devices.** Time-lapse movies were taken using the Nikon Eclipse Ti2 inverted fluorescence microscope integrated with a BSI sCMOS camera and controlled by Nikon Elements software. An oil-immersion 100X object (NA 1.40) was used for imaging. Cells were maintained during imaging at a constant temperature using a microscope live-cell imaging chamber (Haison). Phase-contrast images were acquired at a frame rate of 10 seconds to measure cell length. Cell tracking was performed using custom MATLAB (Mathworks, Natick, MA, USA) software.

**Osmotic force extension assays.** To perform the osmotic-force-extension assay consecutive osmotic shocks were performed using commercial microfluidic plates (BA04, CellASIC, Millipore-Sigma) controlled by the ONIX microfluidic platform. Overnight cells were diluted in fresh media and grown until early exponential phase. Cells were then harvested in the microfluidic chamber at 37°C for 30 min with no shaking to achieve steady-state growth. Prior to the experiments the channels were primed with the desired media for 30 minutes at a pressure of 4psi. The cells were then loaded into the perfusion chamber and osmotic shocks were performed by perfusing media supplemented with the desired sorbitol concentration at a constant pressure of 8 psi. Typically, osmotic shocks were performed every 5 min for a duration of 3 min each. Alexa Fluor 647 dye was added to the media used between the shocks to track media switching. To calculate the change in length resulting from osmotic shock, we determined the intervals of the shocks when the medium was exchanged. Then, for every cell, we calculated the longitudinal strain during each osmotic shock using the formula $\varepsilon_l = (l_f - l_i)/l_i$, where $l_i$ is the length of the cell at the beginning of the interval and $l_f$ is the maximum length of the cell during the interval.

**Super-resolution measurement of single-cell width deformation.** To perform single-cell measurements of width deformation upon osmotic shocks (or enzyme treatment), the cell wall of *B. subtilis* was labelled with fluorescent D-aminoacids (FDDAs, Biotorchis). Before the experiments, cells cultured overnight were diluted in fresh media and cultured until early exponential phase was reached (0.3-0.4 OD). 10mM of RADA was added to the culture 1 hour before the microfluidic experiment. The same concentration of fluorescent D-amino acids was added to the cell loading well. Cultures were back-diluted X100 into the loading well. One image was taken 30 s before the osmotic shock and one image was taken 30 s after the osmotic shock. To track the switching of the media, 0.5mM Alexa Fluor 647 was added to the media used between the shocks.

To quantify changes in width with extreme sub-pixel precision, we developed a fit-free method. Images of labeled single cells pre- and post-shock were computationally aligned by rotating them along a vertical axis. In each image, the portions of the cell containing the septa was cropped, leaving only the cylindrical portion of the cell. Each image was then averaged along the vertical axis to obtain a single-cell average of the fluorescence profile across the width of the cell (Fig. 1C). To calculate circumferential strain without fitting profiles, we normalized the pre- and post-profiles and then scaled one of them along the horizontal (width) axis at sub-pixel intervals, and performed cross-correlation with the other profile.

To measure the circumferential strain versus shock magnitude during dynamic relaxation of cell width (Fig. 4D), we first took a standard curve of width versus time of strain ER476 upon increased induction from 1 mM xylose to 10 mM xylose. We back-diluted an overnight culture induced with 3 mM xylose X100 into LB+1mM xylose. We grew this culture to exponential phase before back-diluting it X100 into LB+10 mM xylose. We then measured cell width on LB agarose pads supplemented with 10 mM xylose every 20 minutes. This confirmed that width began decreasing within 15 minutes after the increase in induction (Fig. S2). Therefore, to measure circumferential strain upon osmotic shock during relaxation, we prepared circumferential osmotic force extension assays at 1 mM xylose induction, as described above, but once we loaded cells into the imaging chamber we perfused them with 10 mM xylose for 15 minutes prior to performing the osmotic shocks. We performed a similar protocol to measure lysis strain during relaxation (see below).

To measure changes in cell width in *C. glutamicum* 0.5mM of RADA was used to label the cells 1 hour before the experiments. For *E. coli* MG1655, the cell wall was labelled with HADA for 1 hour.

**Spp1 endolysin purification.** The bacteriophage endopeptidase, Spp1Lys, was purified by using a previously established protocol[1], from *E. coli* strain CG/pIV::25His. Cells were grown at constant shaking at 28°C until an OD of 0.6-0.8, after that cells were transferred at 42°C for 30 min and at 16°C for 14 hours to induce protein expression. Cells were collected by centrifugation (4000rpm for 30minutes at 4°C) and resuspended in lysis buffer (20mM Hepes, 500mM NaCl, 20mM imidazole, 1% glycerol and 1mM DTT [pH 6.5]). Resuspended cells were lysed by sonication. The supernatant was collected after centrifugation (10000 rpm for 30 min at 4°C) and the enzyme was purified using a column packed with Ni Sepharose High Performance histidine-tagged protein purification resin (Cytiva). The elution buffer was the same as the lysis buffer except that the concentration of imidazole was 500mM. The eluted fraction containing the purified protein was concentrated and the buffer was exchanged to a phosphate-based buffer (50mM phosphate-Na, 500mM NaCl, 25% glycerol, and 1mM DTT [pH6.5]) using a HisTrap Desalting column (Cytiva, 5mL). Whole extract, lysate, and eluted fractions were analyzed by SDS-PAGE and western blot. For western blot, proteins were transferred to a polyvinylidene difluoride membrane using Trans-Blot Turbo system (Biorad). For the detection of SPP1Lys an antibody against the His-tag was used and visualized with a ChemiDoc imaging System. Protein concentrations were determined using a nanodrop (ε= 32.89, M.W.=30.705). The purified enzymes were divided into small aliquots and kept at -80°C in the eluted buffer.

**Hydrolysis assay.** Hydrolysis experiments were performed in microfluidic devices at 37C. Lyophilized powder of lysozyme (Sigma Aldrich) was dissolved in the desired media before the experiments. Purified Spp1 was added directly to the media in the designated chamber at the indicated concentration. Cells were grown in rich media in the growing chamber for 5 minutes prior to the perfusion of the hydrolases. To control for cell lysis, 0.1µM of propidium iodide (Sigma Aldrich) was added to the media with the enzymes. To measure changes in width and length, images were taken 1 minute before and after adding the enzymes, ensuring that cells were not lysed. The measurements were done using custom MATLAB scripts.

**Theoretical phase space.** The theoretical phase space associated with linear elasticity was calculated from Eq. 1. Note that in Eq. 1 the Poisson ratio, n, cannot be defined in the traditional manner since the cell wall is anisotropic. The boundary between the finger trap regime and the no-finger trap regime comes from solving Eq. 1 for the condition $\varepsilon_\theta$=0. The condition that $\alpha^{-1} > \nu^2$ results from substituting the thin shell approximation for the principal surface tensions, $\lambda_l = PR/2$ and $\lambda_\theta = PR$, into Eq. 1 and imposing the physicality condition that the moduli $E_l, E_\theta$, and $E_{l\theta}$ must be greater than zero. Note that this constraint only strictly applies in the limit of the thin shell approximation, which is likely to be approximately true for *B. subtilis* since the thickness of the cell wall is $\approx 40$ nm and the radius of the cell is $\approx 400$ nm. However, contributions of normal stress to the balance of turgor pressure would shift this boundary.

The slices through parameter space that constrain the mechanical properties of the low pressure, finger trap, and stress-softened regimes were found by first finding the slopes of the regression to the circumferential strain versus shock magnitude in the three regimes (slopes $m_q$'s of dotted lines in Fig. 2D), and then substituting these as well as the slope of the regression to longitudinal strain versus shock magnitude, $m_l$ into Eq. 1 as $\varepsilon_{l,\theta} = m_{l,\theta}\Delta C_{out}$. This assumes that in each of these three regimes the cell wall is a linear material near an appropriate reference state. Rather than assuming that the material is linear globally (it obviously is not) this analysis is simply a way to estimate the relative local mechanical properties in the three regimes. Furthermore, all materials are linear for small deformations and non-linear for large deformations, and whether the reference state is the rest state need not affect whether the material behaves locally like a linear elastic material with given parameters $\alpha$ and $\nu$.

**Cell lysis assay.** To measure changes in dimensions in response to lysis, we lysed cells with 5% N-lauroylsarcosine sodium salt. The detergent dissolves the plasma membrane causing the release of the cytoplasmic contents. The cell depleted of the turgor pressure shrinks to its rest length. To measure the change in length in response to lysis, images were taken 1 min before the addition of the detergent and after the detergent was washed away using LB. To calculate changes in dimensions for different levels of Rod complex induction, cells grown overnight in LB with 5mM xylose were back diluted in LB supplemented with the amount of xylose needed to generate cells with the desired diameter (0.5mM, 5mM, and 30mM). The same xylose concentrations were added to the media with detergent and the washing media. The measurements and the calculation of the percentages of change in width and length were done using costume made scripts as described before.

**Pressure measurements.** To measure pressure osmotic force extension experiments were combined with measurements of lysis strains. Pressure was calculated empirically as the hyperosmotic shock magnitude that caused contraction of the cell wall to the rest length observed upon lysis. This was found by finding the intercept of the lysis strain with the regression of the two data points from the osmotic force-extension curve that were immediately above and below the lysis strain. Error in pressure was propagated from the s.e.m. of the lysis strains and the standard error of regression of the osmotic force-extension data points.

**Mathematical model of cell morphogenesis.** To model morphogenesis of a rod-shaped cell where width could change dynamically, we considered a cylindrical cell wall with a constant thickness of 40 nm and pressure of 10 atm. To balance pressure, making the thin-shell approximation, the total longitudinal tension borne by the cell wall is $\lambda_l = \frac{PR}{2}$ and the circumferential tension is $\lambda_\theta = PR$. We discretized the cell wall into 100 layers and allowed the stress and strain to vary across the thickness of the cell wall given these global force-balance constraints. To model dynamic growth and morphogenesis of the cell, we defined an arbitrarily small time step. During each time step we removed one layer (1/100$^{th}$ of wall thickness) from the outer surface of the cell wall and added a unstretched layer to the inner surface of the wall, and then allowed the cell wall to deform to satisfy the global force balance constraints using a standard energy minimization routine.

**Supplementary Figures and Tables**

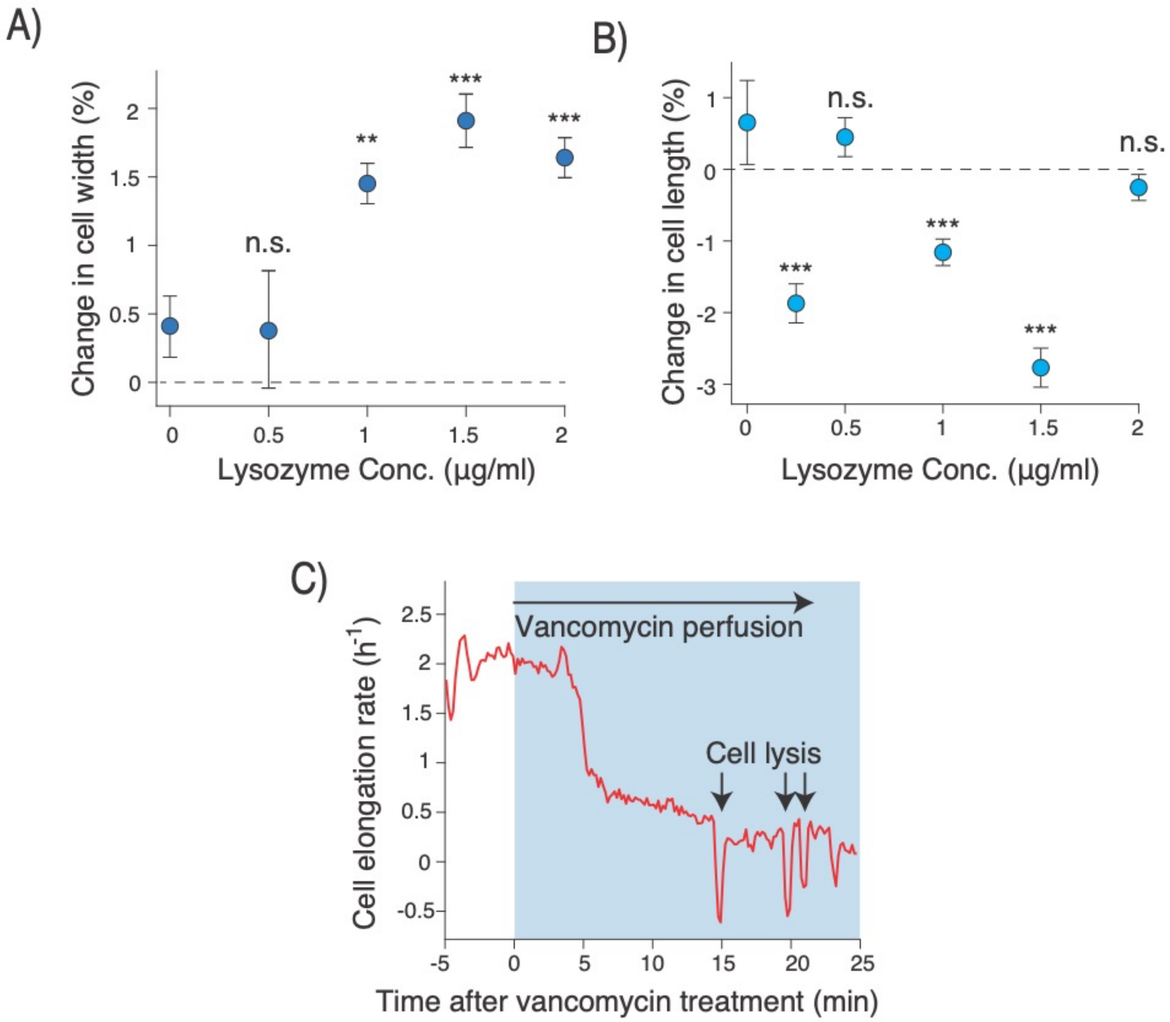


**Figure S1. Lysozyme and vancomycin treatment cause cell widening.** A) Mean change in cell width upon lysozyme treatment. Error bars indicate +/- 1 s.e.m. B) Mean change in cell length upon lysozyme treatment, controlling for cell growth. Error bars indicate +/- 1 s.e.m. C) Population-averaged cellular elongation rate versus time during acute perfusion with 10 mg/mL vancomycin. n=18 cells across 1 experiment.

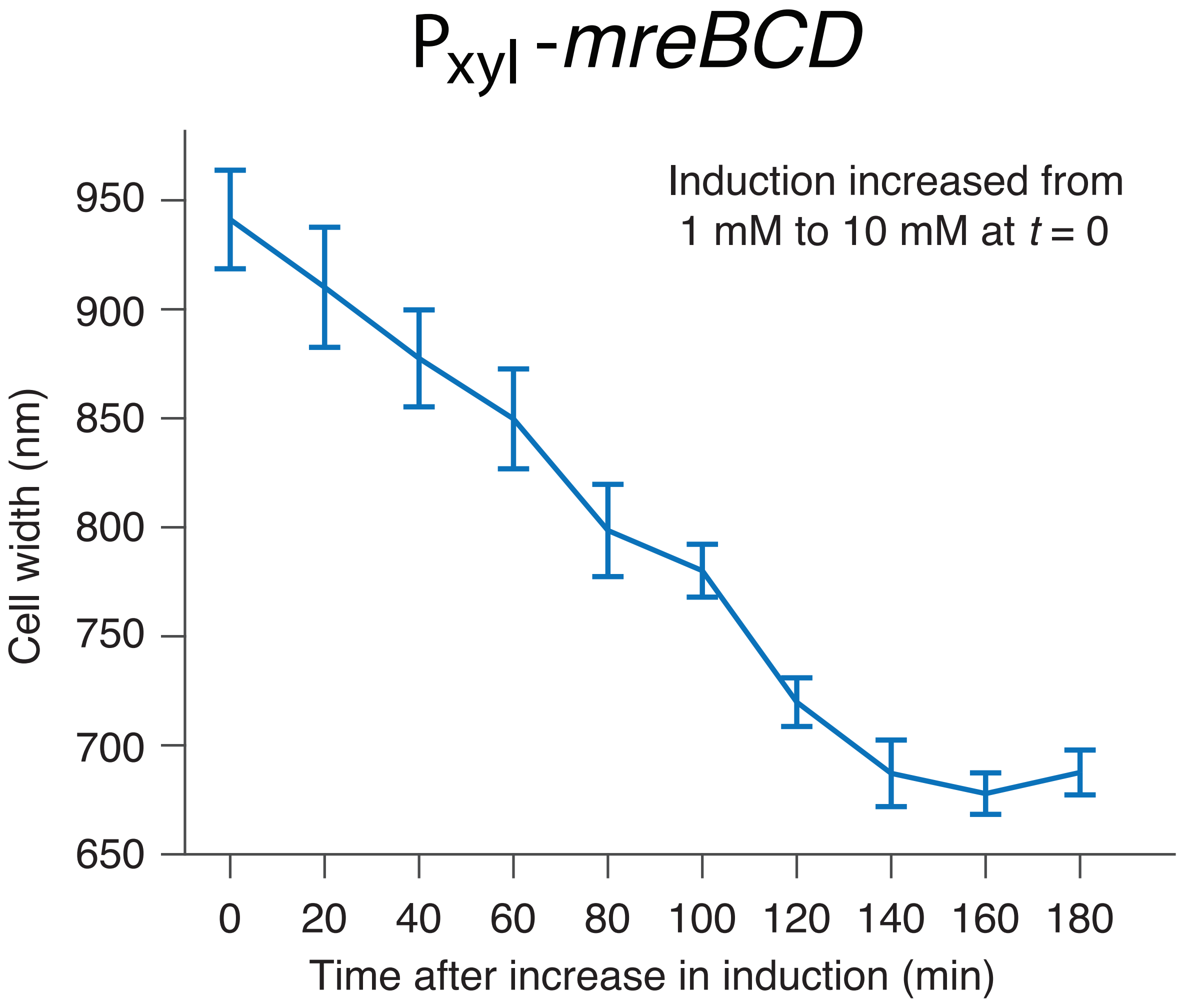


**Figure S2. Acute induction of *mreBCD* causes immediate cell thinning.** Cell width versus time after acute induction of *mreBCD*. Cells were grown to exponential phase in 1 mM xylose and then xylose concentration was increased to 10 mM. n=20 cells across 1 experiment for each time point.

| Strain or Plasmid | Genotype | Relevant features | Source/Reference |
|---|---|---|---|
| *B. subtilis* PY79 | wild-type | | Lab stock |
| bMD545 | *PY79 amyE::erm Pxyl-mreBCD, ΔmreBCD::spc PmreB-minCD* | Xylose inducible induction of mreBCD operon | Garner Lab[2] |
| pPB001 | pIV::25His | pIVEX2.3d derivative carrying SPP1 endolysin gene 25 | São-José Lab[1] |
| *C. glutamicum* | wild-type | | Theriot Lab |
| *E. coli* | wild-type MG1655 | | Lab stock |

**Table S1. Strains and plasmids used in this study.**
